# Connectivity Is All You Need: Inferring Neuronal Types with NTAC

**DOI:** 10.1101/2025.06.11.659184

**Authors:** Gregory Schwartzman, Ben Jourdan, David García-Soriano, Arie Matsliah

**Author notes:** These authors contributed equally (Gregory Schwartzman, Ben Jourdan).

## Abstract

Recent advances in electron microscopy and computer vision have enabled the mapping of complete wiring diagrams—called connectomes—of brain regions and even entire brains. The emergence of these increasingly large-scale connectomic datasets have intensified the need for efficient and accurate neuronal cell type identification. Traditional approaches rely on labor-intensive analyses of molecular, anatomical, and physiological features. As a step toward fully automated neuronal cell type classification, we present **NTAC** (Neuronal Type Assignment from Connectivity) — a method for grouping neurons into cell types based solely on synaptic connectivity. Our approach is grounded in the hypothesis that synaptic connectivity is key to determining neuronal cell types (Seung 2012), and our results provide the strongest evidence to date supporting its validity.

NTAC comes in two flavors: a **semi-supervised** variant that requires some of the neurons to be labeled, and an **unsupervised** variant that requires no labels at all. The first can be naturally formalized as a learning problem on graphs: given labels for a subset of neurons, use connectivity to label the rest. We present a simple and fast algorithm that achieves over 95% accuracy on the fly’s visual system starting with just 2% of neurons labeled, within minutes on a standard PC. In contrast, morphology based parallel achieves lower accuracy even on smaller datasets and with significantly more labels.

Formalizing the problem as a fully unsupervised learning task is more challenging. To address this, we introduce a novel computational problem, *approximate equitable partitioning*, and design an efficient heuristic for it, using the semi-supervised algorithm as a subroutine. Remarkably, the unsupervised method achieves ∼70% accuracy on the fly’s visual system, drastically outperforming its morphology based parallel and providing concrete evidence that synaptic connectivity alone can reveal cell types.

Both variants of NTAC are evaluated on multiple state of the art connectomes including optic lobes, central brain and nerve cord of adult fruit flies.

## Introduction

Connectome reconstruction involves mapping the network of neurons and synaptic connections within the brain, typically through electron microscopy (EM) imaging, AI-assisted segmentation and synapse detection, followed by human expert proofreading and annotations leading to classification of cell types^1^. Cell types are the fundamental units for interpreting circuit function and comparing brain organization across species. In increasingly large connectomic datasets, cell types provide the necessary level of abstraction—making otherwise intractable networks understandable by collapsing complexity into interpretable units.

When new connectomes are reconstructed, there are two potential approaches for cell typing. If a fully annotated connectome of the same species (and brain region) is available, the cell types can be mapped to the new dataset using morphology and connectivity based neuron-to-neuron comparisons. Specifically for *Drosophila melanogaster* (fruit fly, a popular model organism), one of the primary tools used for morphological clustering of neurons is NBLAST (Neuron BLAST) (Costa et al. 2016), which compares pairs of neurons based on their shape and position. While this process is typically not fully automated and requires expert supervision and corrections, it has been recently carried out on multiple datasets to accurately recover cell types in the central brain (Schlegel et al. 2024; Winding et al. 2023).

In the second scenario, when a connectome is typed from scratch (for example, new species or new parts of the nervous system that were never typed before), cell typing becomes more challenging. For *Drosophila* **central brain** (where cells of same type are co-located and morphologically stereotyped) clustering approaches are typically invoked to define cell types, using NBLAST first and refined with connectivity analysis - see the methods described in (Schlegel et al. 2024; Scheffer et al. 2020; Matsliah et al. 2024; Nern et al. 2024; Winding et al. 2023). However, morphology-based clustering lacks robustness in numerous neuronal types in the optic lobes—where the majority of the neurons are located in small model organisms like *Drosophila* (Dorkenwald et al. 2024; Matsliah et al. 2024; Nern et al. 2024) —due to the diverse morphologies and spatial positions of same-type neurons that tile the visual field.

Furthermore, partial reconstructions of mammalian brain connectomes provide evidence of numerous cell types spread across large regions in a similar “tiling” pattern (Bae et al. 2018; Schneider-Mizell et al. 2025).

### Overview of NTAC

NTAC is an automated method for detecting neuronal cell types relying solely on connectivity, aiming to provide a more robust and efficient approach to cell type classification in new large-scale connectomic datasets. NTAC can function in two modes: an unsupervised mode (**unseeded** regime) requiring only raw connectivity data, and a semi-supervised mode (**seeded** regime) where a small fraction of labeled neurons guides classification.

We demonstrate its accuracy using the cell types from state of the art connectomic datasets: FlyWire Consortium’s female *Drosophila* central brain and optic lobes (Dorkenwald et al. 2024; Schlegel et al. 2024; Matsliah et al. 2024) and the Janelia Research Campus’s male *Drosophila* visual system (Optic Lobe) (Nern et al. 2024) and ventral nerve cord (MANC) (Takemura et al. 2024). In the seeded regime, where known cell types are provided for a **small percentage** of nodes, the accuracy approaches 100% on the visual systems (where the types contain many cells^2^ and morphology-based methods perform poorly), and exceeds 90% on the full brain. By contrast, applying a k-nearest neighbor (k-NN) classifier to morphological similarity scores (e.g., NBLAST) performs substantially worse on all datasets, except the central brain. When considering top-5 accuracy, NTAC exceeds 90% across **all brain regions**, making it an effective tool for human-assisted labeling even in regions where it does not excel.

In the fully unsupervised “unseeded” regime, the accuracy of NTAC nears 70% on the visual systems. By contrast, standard clustering algorithms to morphological similarity scores (e.g., NBLAST) performs substantially worse, resulting in accuracy less than 10% (see Fig. 9). For the full brain, a substantially harder case with thousands of unique cell types, NTAC still achieves 52% accuracy. To the best of our knowledge, this is the first **unsupervised** algorithm to produce meaningful results for the cell type classification problem.

These results highlight the practicality of connectivity-based inference of cell types in large scale connectomic datasets, and provide concrete evidence that synaptic connectivity alone contains enough information for identifying neuronal cell types.

NTAC implementation is available as an open-source Python library, optimized for optional CUDA/GPU acceleration in the unseeded version. On a full fruit fly brain dataset, it converges within minutes in seeded mode and within a few hours in unseeded mode on a modern PC (see the Code Availability and Evaluation sections).

**Figure 1.**
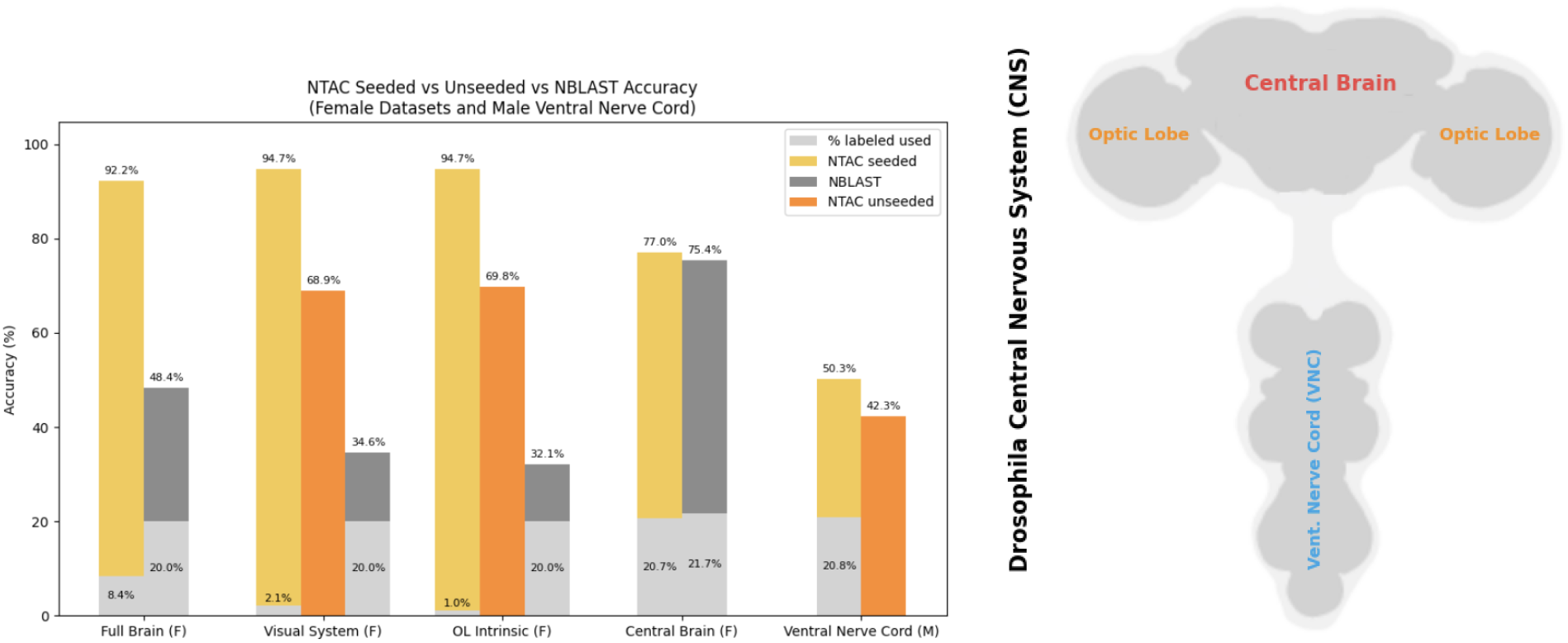
Summary of selected results – we omit those for the male visual system for brevity, as they outperform the female brain (see Evaluation section for full results). F/M denote Female and Male Drosophila connectomic datasets. Full brain contains both the Central Brain and Visual System (which includes neurons in the optic lobe as well as projection neurons to/from the optic lobes and central brain). OL Intrinsic is a subset of the visual system containing neurons that are intrinsic to the optic lobes (all their synapses are within the optic lobes). Ventral Nerve Cord is separate from all other regions, together with Full Brain comprising the entire CNS (central nervous system) of the fly. For the “Visual System”, “OL Intrinsic” and “Central Brain” the results are for both sides of the brain. Percentage of labeled cells used in the seeded regime depends on the region - for example, in the optic lobes 1% labels suffice to have one cell of each type labeled since there are >100 cells per type, while in the ventral nerve cord approximately 20% are required since there are 4.5 cells per type on average and at least one example is labeled per type. Due to the large number of types, our unseeded algorithm requires prohibitively long runtimes on the central brain and full brain datasets; hence, these results are omitted.

**Figure 2.**
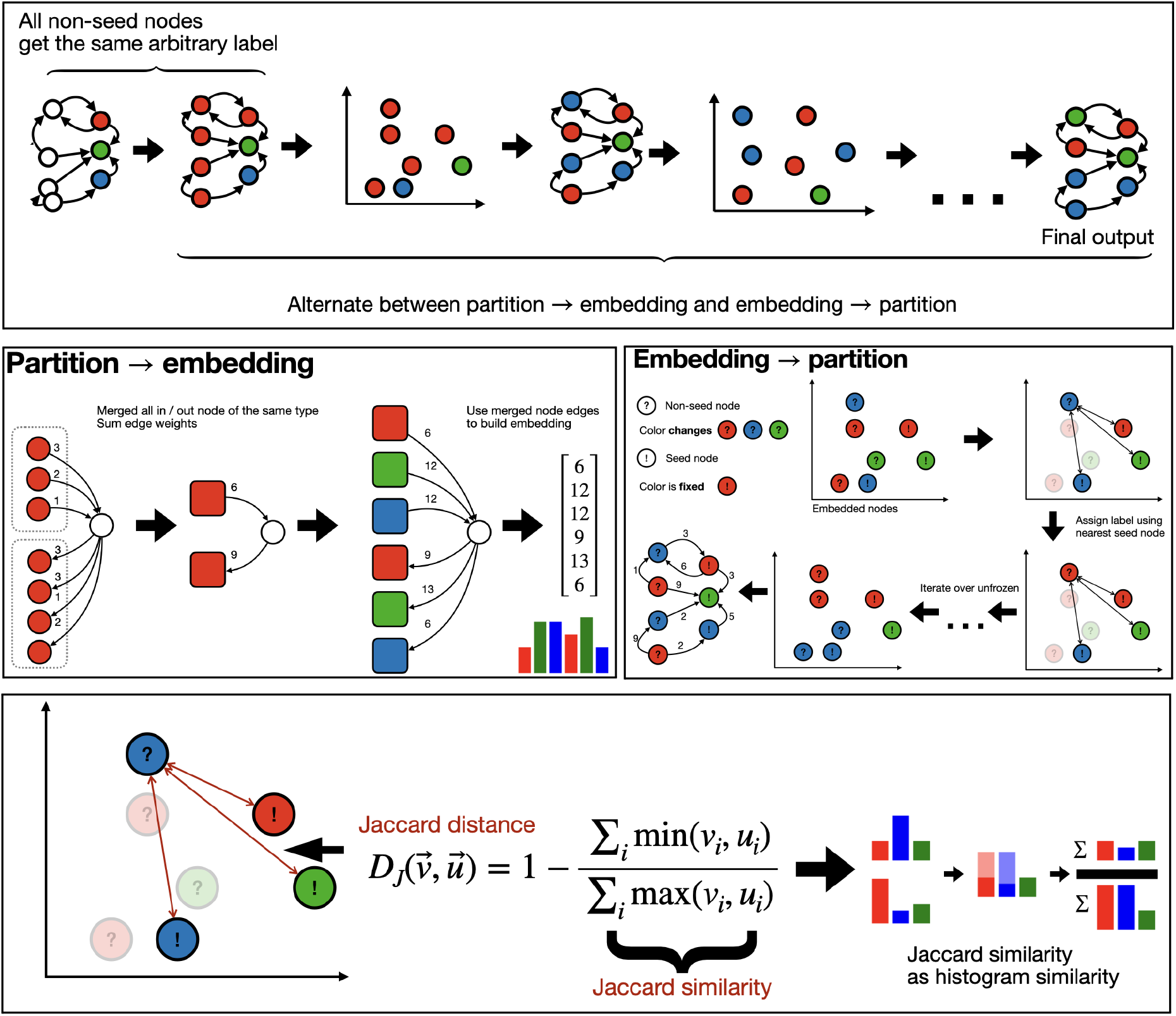
Visual illustration of the main components of the seeded algorithm. Top: overview of the entire algorithm. Middle: illustration of how an embedding is constructed from a partition and vice versa. Bottom: illustration of nearest neighbors via Jaccard distance.

## Related work

### Semi-supervised approaches

Seeded NTAC takes as input a neuronal wiring diagram with a small subset of nodes labeled by cell type, and infers labels for the remaining, unlabeled nodes using graph-based analysis of connectivity patterns. In the machine learning setting, the seeded variant of our problem corresponds to a transductive learning problem on graphs (Kipf and Welling 2017). There are many datasets considered in the machine learning literature, such as citation networks, protein networks, fraud detection, and so on. To the best of our knowledge, all current state of the art results rely on Graph Neural Networks (GNNs) (Kipf and Welling 2017).

Seeded NTAC falls within the iterative classification framework (Sen et al. 2008), a predecessor to current deep learning methods. It does not require any node features (unlike most GNNs), and, at its core, relies on a simple nearest neighbor classifier under a changing distance metric which is tailored to connectome data. This simplicity allows for inherent parallelization, resulting in a fast running time.

Several deep learning frameworks have been proposed for neuronal typing, with reported accuracies often exceeding 90% (Troidl et al. 2025),(Liao, Wan, and Du 2024),(Xiong et al. 2024),(Hanbo Chen, Jiawei Yang, Daniel Iascone, Lijuan Liu, Lei He, Hanchuan Peng, Jianhua Yao, n.d.), (Jiang et al. 2023). However, these methods either operate on small datasets with few types or deliberately exclude rare cell types—an approach that does not generalize to anatomically complex regions such as the central brain or ventral nerve cord. Moreover, they rely on dense supervision (e.g., 90% of neurons labeled), whereas NTAC achieves competitive performance using as little as **1% labeled data**. Most importantly, all previous works rely on morphological data while NTAC does not.

Particularly relevant to this work are the results of (Troidl et al. 2025), and (Liao, Wan, and Du 2024). Troidl et al.’s Point Affinity Transformer classifies neurons based solely on morphological shape, embedding them as sparse 3D point clouds and utilizing a k-NN classifier. They report 81% accuracy on predicting cell type families (superclasses of fine-grained cell types which NTAC predicts) in the FlyWire optic lobe and 77% accuracy on the Hemibrain dataset (a subset of our “central brain” dataset) - both with types containing less than 40 neurons excluded. The deep learning framework in (Liao, Wan, and Du 2024) takes as input both neuron skeleton data and the full connectome, reporting 91.7% accuracy on Hemibrain (reduced to 191 types) and 93.6% on the H01 human cortical dataset. Both works use around 90% of the data for training. NTAC achieves near perfect results on the optic lobe using only a small fraction of labeled nodes without any type reduction. While we cannot make a direct comparison with the modified Hemibrain datasets of (Troidl et al. 2025) and (Liao, Wan, and Du 2024), we note that using the full 4000+ types of the central brain, NTAC achieves 87% accuracy when using 80% labeled nodes in just 15 seconds - orders of magnitude faster than (Liao, Wan, and Du 2024). This speed discrepancy is inherent as the connectome data is much easier to process compared to morphological data ((Liao, Wan, and Du 2024) note that their main bottleneck is running the morphological data through the classifier).

### Unsupervised approaches

Our unsupervised problem, approximate equitable partitioning, is a relaxation of equitable partitioning (Schwenk 1974), where nodes in the same cluster have identical degree counts to every other cluster (see “Algorithm Description”). Such a partition into a minimum number of clusters may be found in polynomial time. However, this rigid definition often yields partitions with one cluster per node, rendering them uninformative for noisy biological data. While equitable partitions are well-studied in spectral theory (Godsil 2017), approximations capturing near-equivalence have not been explored, to the best of our knowledge.

Other conceptually related graph analysis methods such as graph summarization (Riondato, García-Soriano, and Bonchi 2017) create a concise, lossy representation by grouping vertices into “supernodes” and collapsing edges, optimizing for global accuracy of individual edge information. However, a perfectly equitable partition might incur high reconstruction cost in that sense. Szemerédi’s regularity lemma (Riondato, García-Soriano, and Bonchi 2017; Szemerédi 1975) and its algorithmic forms (Schwenk 1974; Alon et al. 1994; Frieze and Kannan 1999) partition graphs into pseudorandom blocks, controlling aggregate edge counts over large enough subsets of every pair of clusters. In contrast, our method only considers per-node edge counts to complete clusters, making the two notions incomparable.

Unseeded NTAC uses a strategy akin to Gonzalez’s k-center algorithm (Chierichetti et al. 2010; Gonzalez 1985), iteratively selecting distant centers. But since the connectivity-based representations of nodes depend on the partition, we rely on *seeded* NTAC as a subroutine to determine the best partition given only a set of center nodes.

## Algorithm Description

To present NTAC we begin by describing the seeded algorithm followed by the unseeded version that uses the seeded variant as a subroutine.

**Notation**. Let *G*(*V, E, w*) denote the directed weighted input graph, where *w*: *E* → *R*^+^. For *A, B*⊆ *V*, denote by *e*(*A, B*) = ∑_*u*∈*A, v*∈*B*_*w*((*a, b*)) the weighted number of edges from *A* to *B*. Similarly, for *u*∈*V*, define *e*(*u, B*) = *e*({*u*}, *B*) (the out-degree of *u* w.r.t *B*) and *e*(*A, u*) = *e*(*A*, {*u*}) (the in-degree of *u* w.r.t. *A*). Finally, for an integer k let [k] = {1, 2, …, k}.

### NTAC - Seeded

The input for seeded NTAC is a weighted directed graph with partial node labels. Nodes correspond to neurons, edges to synaptic connections, edges weights to the number of synapses per connection and node labels are neuronal cell types. NTAC assigns labels to all unlabeled nodes such that the formed labeling approximately recovers neuronal cell types in the corresponding connectome.

Let us refer to the set of labeled nodes as seeds, and to the set of unlabeled nodes as non-seeds. Let us assume we have k unique labels in the range [k] = {1,…,k}.

Given the input graph and partial node labels, the algorithm begins by assigning all non-seed nodes to the same arbitrarily chosen label. This guarantees that all nodes are labelled. The algorithm then repeats the following steps for a given number of iterations:

1. Compute node embeddings using the current labeling
2. Assign new node labels using the embeddings

A node embedding is a mapping from nodes to vectors. In our case the embedding has 2k dimensions. Given the current labelling, the first k dimensions of the embedding correspond to the total incoming weight to the node from each label: the i-th entry in the embedding vector for node v is the sum of the weights of incoming edges that point to v from nodes with label i. Similarly, the last k dimensions correspond to the total outgoing weight from the node to each label: the (k+i)-th entry corresponds to the sum of the weights outgoing to label i. More formally:

**Definition (degree-count embedding)**

Given a *k*-partition *𝒫* = {*C*_1_, …, *C*_*k*_} of *V* and a node *u*∈*V*, the *degree-count embedding (embedding* for short) of *u* is the vector *f*(*u*) = *a*∈ℝ^2*k*^ defined by: *a*[*j*] = *e*(*u, C* _*j*_) and *a*[*j* + *k*] = *e*(*C*_*j*_, *u*), for *j* ∈ [*k*].

After constructing the embeddings, we update the labels of each non-seed node by assigning each node to the seed with the closest feature vector. Closeness is measured by *Jaccard distance* (one minus *Jaccard similarity*), defined as

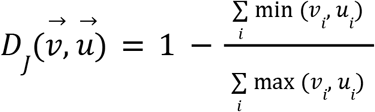

where 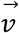 and 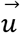 are embedding vectors of the corresponding nodes. Intuitively, this measures the distance of the neighborhood between nodes, with respect to the current labeling. Pseudocode for our algorithm is given below:

#### Algorithm 1

Seeded NTAC

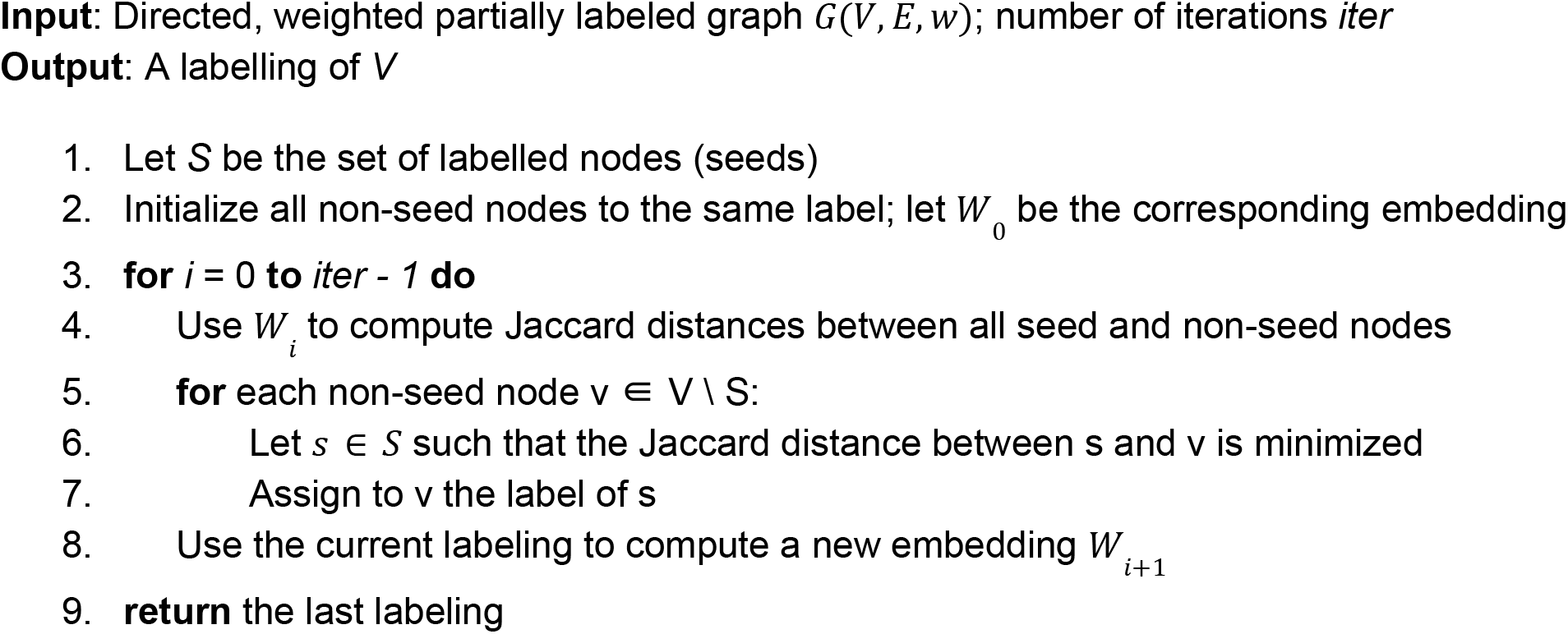

#### Returning Top-k labels

Seeded NTAC can also return a ranked list of labels for each node, along with corresponding similarity scores. For every node v, we compute the Jaccard distance to all seed nodes, sort them by increasing distance, and extract the top-k closest *unique* labels.

### NTAC - Unseeded

We present an unsupervised version of NTAC requiring no labels as input. Given a directed, weighted graph *G* = (*V, E, w*) and a target number of types *k*, our method incrementally constructs a partition of *V* into *k* clusters.

#### Problem formulation - approximate equitable partitioning

We need to capture the notion that cells of the same type share a common connectivity pattern. Given a partition, the connectivity pattern of a single vertex *v* is described by its degree-count embedding *f(v)* from the previous section, but here we would like to be able to speak about the connectivity pattern of an entire cluster. To this end, let M = (*M*_1_, …, *M*_*k*_)∈ℝ^2*k*^ denote a sequence of vectors, which play the role of centroids of the embeddings of each cluster in a given partition. Given a k-partition *𝒫* and M as above, we define the following *Jaccard cost*:

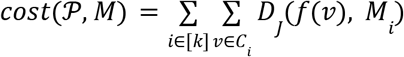

which measures closeness (in Jaccard distance) between the embeddings of nodes and the centroids (elements of M) of their assigned clusters.

We define the following problem:

**Problem (approximate equitable partitioning)**

Given a directed weighted graph *G* and a number of clusters *k*, find a partition *𝒫* into at most *k* non-empty clusters minimizing

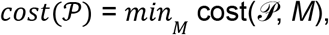

where *M* ranges over all *size-k* sequences of 2*k*-dimensional vectors *M* = (*m*_1_, …, *m*_*k*_).

Observe that, given a partition *𝒫*, the optimal choice for each *M* is the vector minimizing

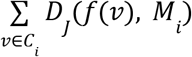. By definition, this is the so-called Jaccard median of the embeddings of *C*_*i*_.

While the Jaccard median is NP-hard to compute, it admits a polynomial-time approximation scheme (Chierichetti et al. 2010). In practice, our implementation uses coordinate-wise medians (or, if desired, coordinate-wise means) of the embeddings of *C*_*i*_.

#### Overview

A key technical challenge is that the *embeddings depend on the partition itself*, creating a circular dependency. This rules out standard clustering approaches which assume fixed feature vectors. To address this, we first consider the simple special case when there is a solution *𝒫* of cost 0. This occurs when *G* has an *equitable partition* into *k* clusters. A standard result from graph theory (Chierichetti et al. 2010; Gonzalez 1985; Schwenk 1974) gives a simple algorithm to find the equitable partition into a minimum number of clusters (which is unique) by iterative refinements:

- Start with the trivial partition *𝒫* into one cluster.
- If all *𝒫*-embeddings are equal, return *𝒫*.
- Otherwise, create a partition *𝒫*’ by putting two nodes in the same cluster if and only if they share exactly the same embeddings relative to *𝒫*.
- Set *𝒫*’ = *𝒫* and go back to step 2.

In practice, due to biological variability and reconstruction errors, an exact equitable partition for the neuronal connectivity graph will necessitate a large number of clusters, nearly equal to the total number of neurons. Hence this does not provide a viable solution.The key challenge in adapting the classical algorithm to the approximate setting lies in determining when two embeddings from cells currently assigned the same type are different enough to warrant separating them into distinct clusters.

Our proposed heuristic builds a sequence of *seed vertices s*_1_, …, *s* _*k*_, one at a time. On each iteration *l*, we use the embeddings of the current partition to select greedily an additional seed *s* _*l*+1_ which can serve as the center of a new cluster. Then we compute the new partition into *l* + 1 clusters using the seeded NTAC algorithm from the previous section. Finally, we recompute the current set of *l* + 1 seeds so that they better reflect the centers of each cluster found. The algorithm stops when the desired number *k* of clusters is reached. Details follow.

#### Initialization

The first partition *𝒫*_1_ is the trivial partition into one cluster. We compute for each node *u* ∈ *V* an embedding *f*(*u*) = [*e*(*u,V*), *e*(*V,u*)] capturing its in and out degrees. The first seed *s*_1_ is selected as the vertex whose embedding is closest (in Jaccard distance) to the coordinate-wise median of all embeddings.

#### Adding a new seed

At each step *l*, given *l* < *k* seeds and a partition *𝒫*_*l*_ into *l* clusters, we recompute feature vectors relative to the current partition, and for each cluster *i*∈*[l]*, a median embedding *M* _*i*_. Then we score each non-seed by how much it would reduce total Jaccard cost if it formed a new cluster along with the vertices that are closer to it than to their current medians (holding other cluster medians fixed). Formally,

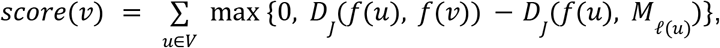

where *ℓ*(*u*) represents the current cluster assignment of *u*_*l*+1_. The next seed *s* is chosen as the non-seed vertex with maximum score.

#### Computing a new partition

After selecting the new seed, we run the seeded NTAC subroutine, obtaining an updated partition *𝒫*_*l*+1_ with one more cluster.

#### Updating the seeds

The next step is to recompute the set of seeds, such that the embedding of each seed is closest to the median of each current cluster. The result is a new set of seeds *𝒮*_*l*+1_. Note that the choice of *s*_1_ during initialization was a special case of this.

#### Termination

The process repeats until *k* seeds have been selected, yielding the final partition.

##### Algorithm 2

Unseeded NTAC

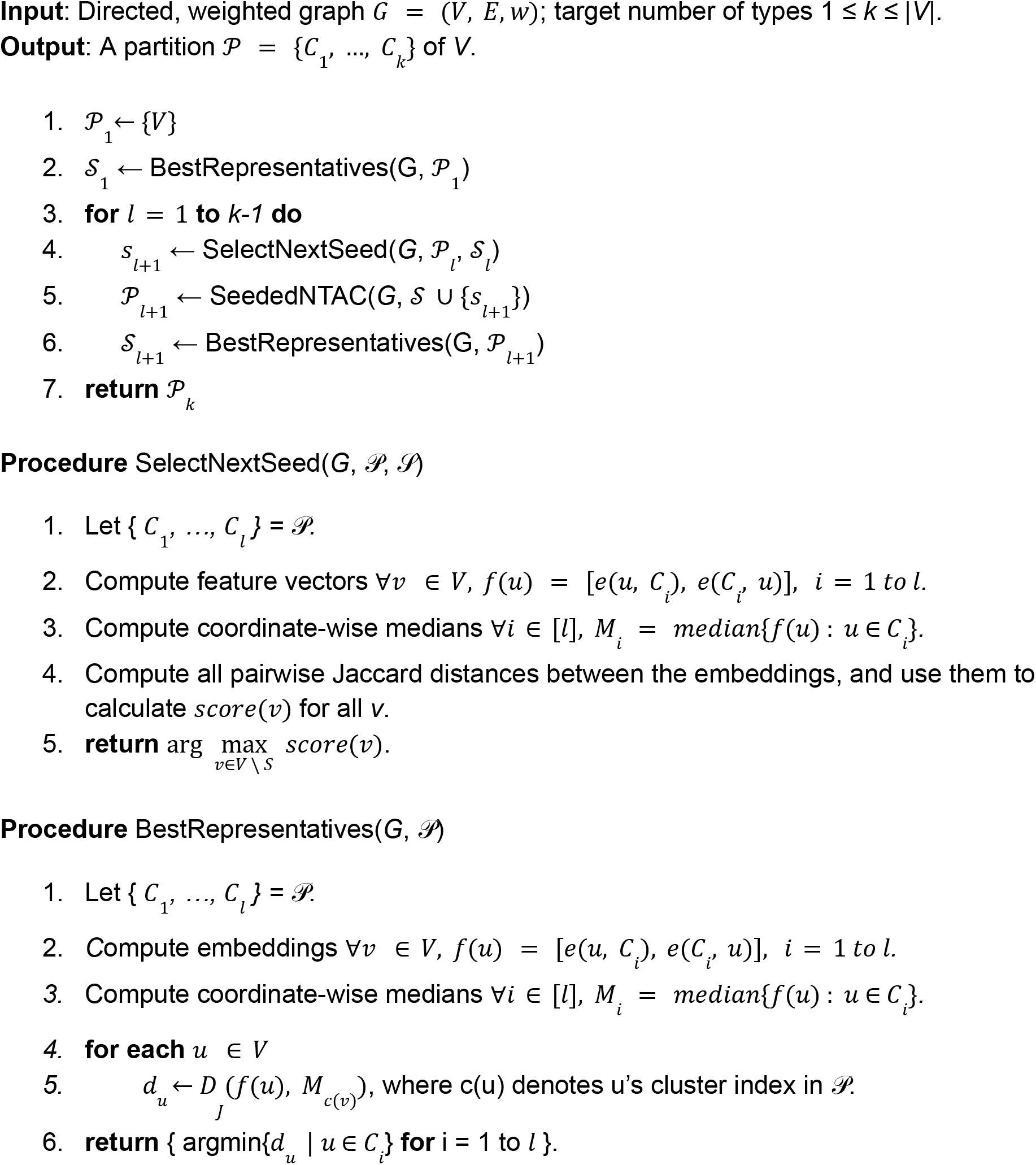

#### Computational complexity

On a graph with *n* nodes and *m* edges, if R iterations are used by SeededNTAC, the overall runtime of UnseededNTAC is *O*((*n k*)^2^ + *R* (*n k*^3^ + *m k*)). For small R (the typical case as SeededNTAC converges quickly), this simplifies to *O*((*n k*)^2^). The computational bottleneck is computing pairwise Jaccard distances within SelectNextSeed.

Let us first analyze SelectNextSeed. Computing embeddings for an *l*-partition (lines 1-2) takes linear time, *O*(*n* + *m*). The coordinate-wise medians of the 2*l*-dimensional embeddings of a size-*c* cluster can be computed in time *O*(*cl*), hence all coordinate-wise medians (line 3) can be computed in *O*(*nl*). Jaccard distance calculations from all *n* embeddings to a set of size *s* take time *O*(*nsl*). Taking *s=n*, we see that lines 4-5 runs in *O*(*n*^2^*l*), which is then the overall complexity of SelectNextSeed.

As for BestRepresentatives, we have seen that lines 1-3 can be run in *O*(*nl* + *m*). The main loop takes *O*(*nl*), so its complexity is *O*(*nl* + *m*).

Finally, let us consider the main algorithm UnseededNTAC. Initialization (lines 1-2) is dominated by BestRepresentatives with *l* = 1, and hence runs in *O*(*n* + *m*). Each iteration of the main loops takes *O*(*n*^2^ *k*) for SelectNextSeed, *O*(*R* (*n k*^2^ + *m*)) for SeededNTAC, and *O*(*nk* + *m*) for BestRepresentatives. Thus, the complexity of each of the *k* iterations is *O*(*n*^2^ *k* + *R* (*n k*^2^ + *m*)).

#### An alternative: return the best partition

Alternatively, UnseededNTAC could be made to return the partition with minimum *cost*(*𝒫* _*i*_), among the *k* partitions found, rather than the last partition found. Our experiments suggest this is often a better choice in practice, as forcing the addition of more clusters may sometimes increase cost and reduce accuracy.

#### Practical speedup

The practical running time is dominated by seed selection, and in particular by the need to compute all pairwise Jaccard distances. Although this may be easily parallelized, for a further speedup we have implemented an optional parameter *T*∈(0, 1) which is used to evaluate only a *T*-fraction of all nodes as potential seeds within SelectNextSeed, thus reducing its cost at a negligible loss in accuracy. To determine the set of *Tn* potential seeds, we pick, for each cluster, the node farthest away from its median in Jaccard distance; then the second-farthest within each cluster; and so on, until reaching a set Q of *Tn* potential seeds. Finally, we return arg 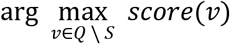 in place of the last line of SelectNextSeed.

## Evaluation

### Seeded NTAC

We applied our method to the state of the art *Drosophila* connectomes: FlyWire full-brain connectome (Dorkenwald et al. 2024) and the Janelia Research Campus’s optic lobe (Nern et al. 2024) and ventral nerve cord (Takemura et al. 2024) connectomes. The results demonstrated that our approach accurately classified neurons into known cell types. First, we present our accuracy scores across different regions of the brain for the FlyWire dataset as a function of percentage of correctly labeled (seed) neurons. We observe that NTAC achieves near perfect accuracy on the optic lobes, and even for the central brain (which consists of mostly cell types with very few cells), we achieve high accuracy. Note that the plot starts from a rather high percentage of labeled nodes. This is due to our assumption that we have at least one labeled node per class and the fact that the central brain contains many cell types. We can make do with far fewer labeled nodes for specific brain regions.

**Figure 3.**
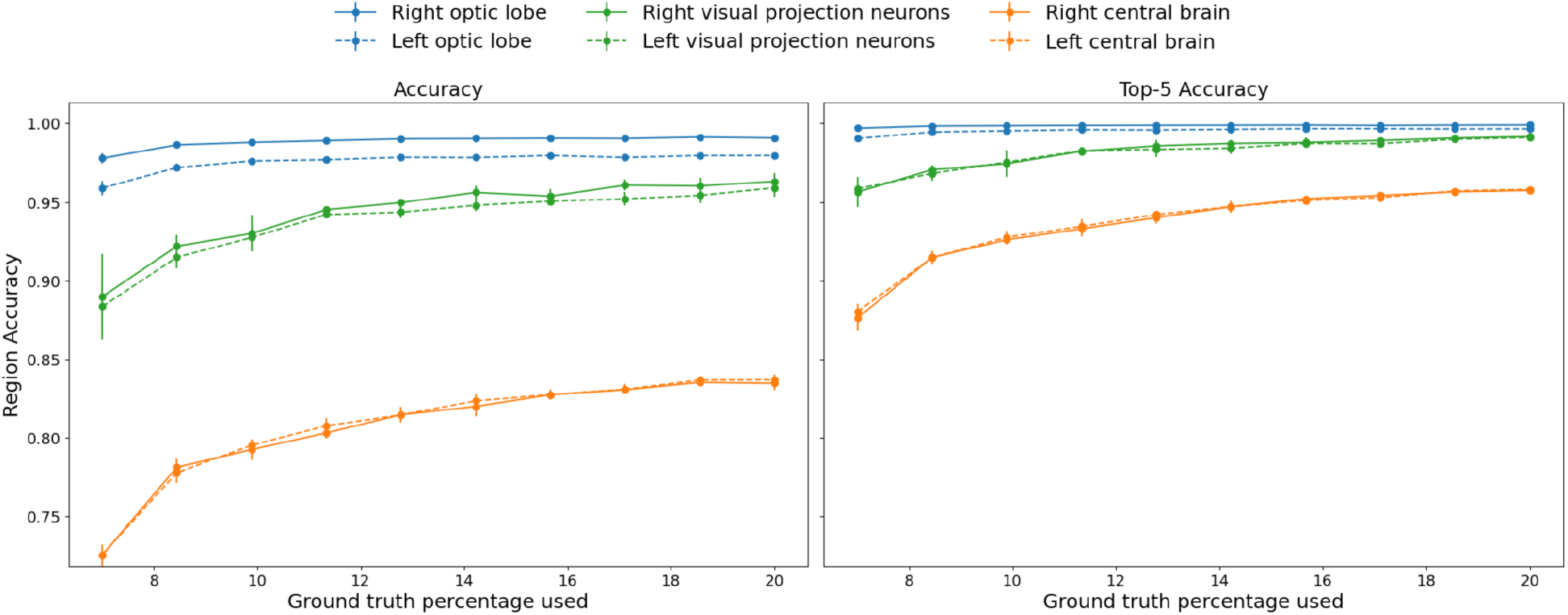
Accuracy and Top-5 accuracy curves for different brain regions as a function of % labeled nodes provided to NTAC. We guarantee that there is at least one labeled node per class.

**Figure 4.**
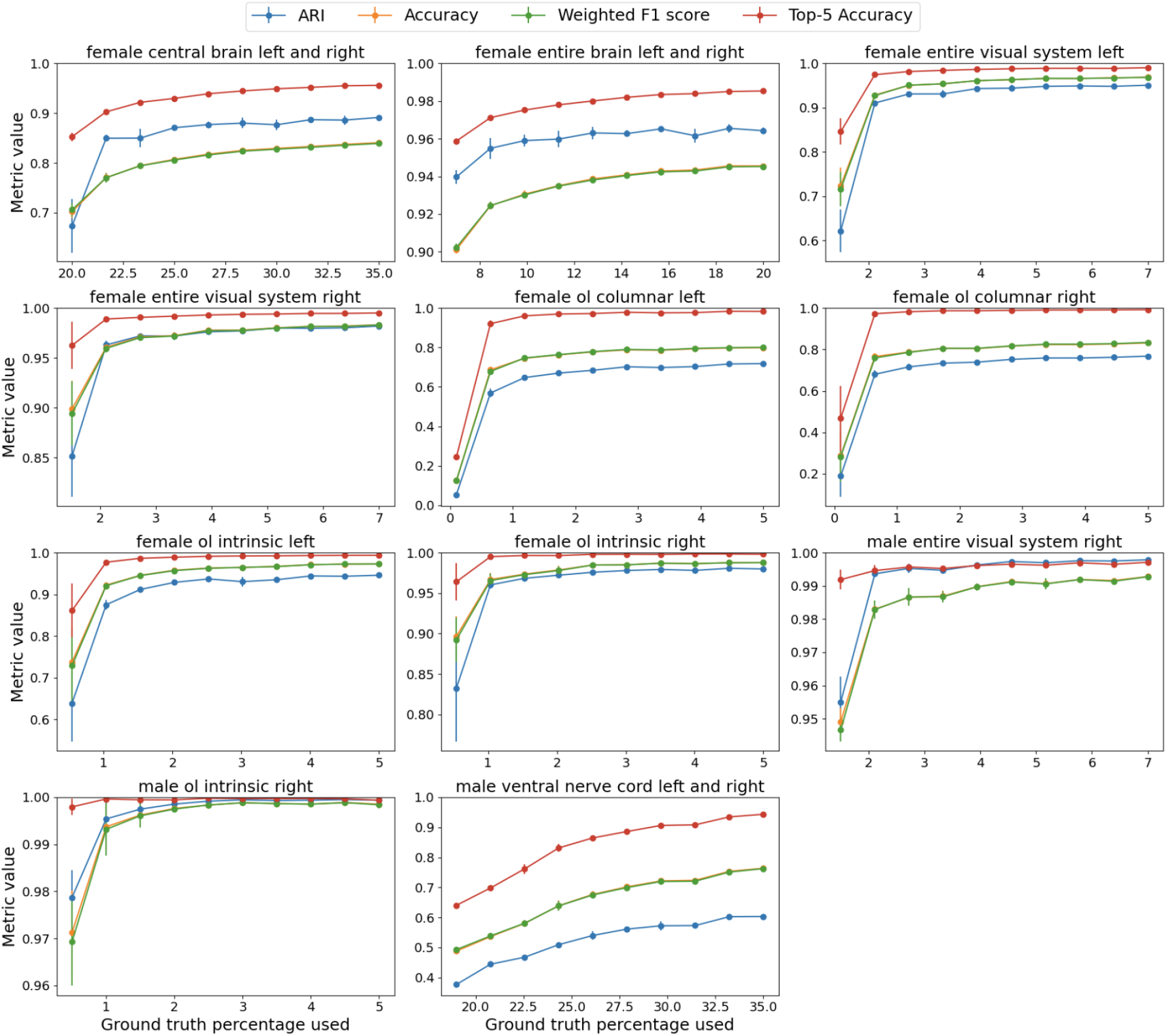
Detailed plots per dataset. We plot accuracy, top-5 accuracy, ARI and weighted F1 as a function of labeled nodes. We guarantee that there is at least one labeled node per class. Note the low accuracy on the OL Columnar dataset—in the Supplemental Data section, we show that this is due to several cell types being indistinguishable based on connectivity alone in this limited dataset, despite being separable in larger datasets.

**Figure 5.**
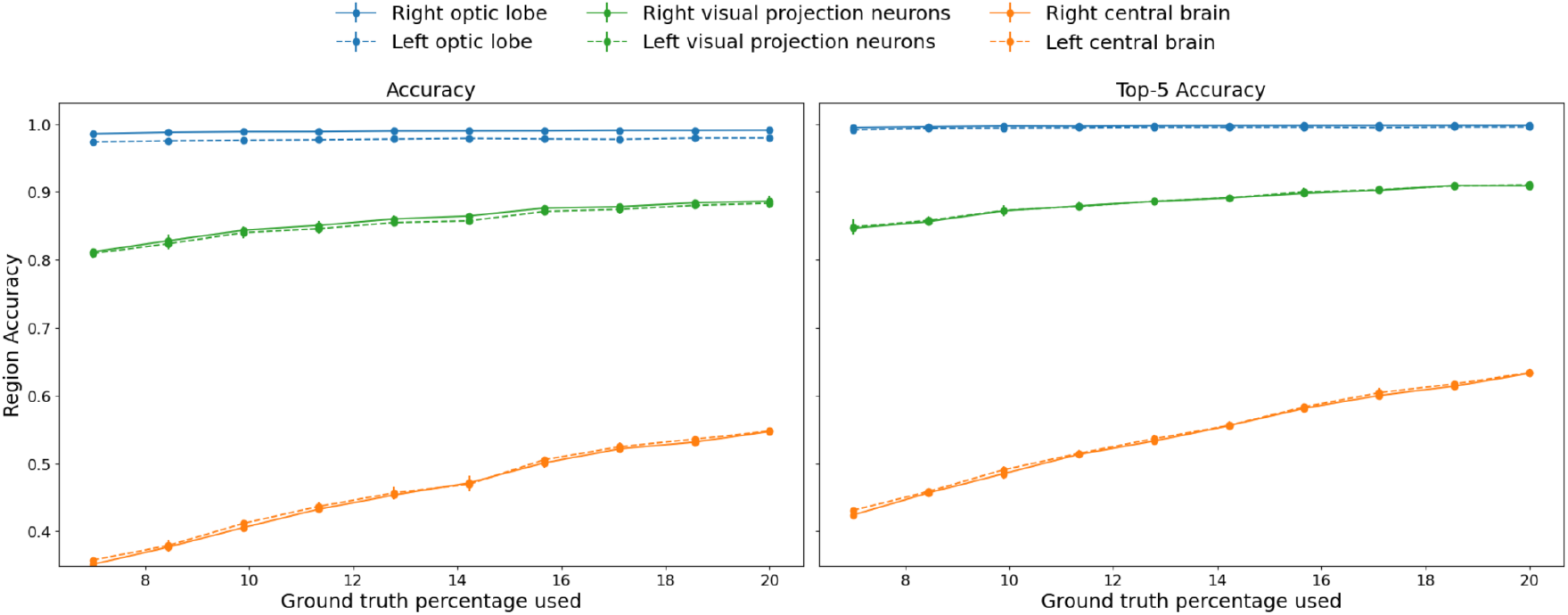
Accuracy and top-5 accuracy curves for different brain regions as a function of %labeled nodes provided to NTAC. We sample the labeled nodes uniformly at random, and do not guarantee that there is at least one labeled node per class.

**Figure 6.**
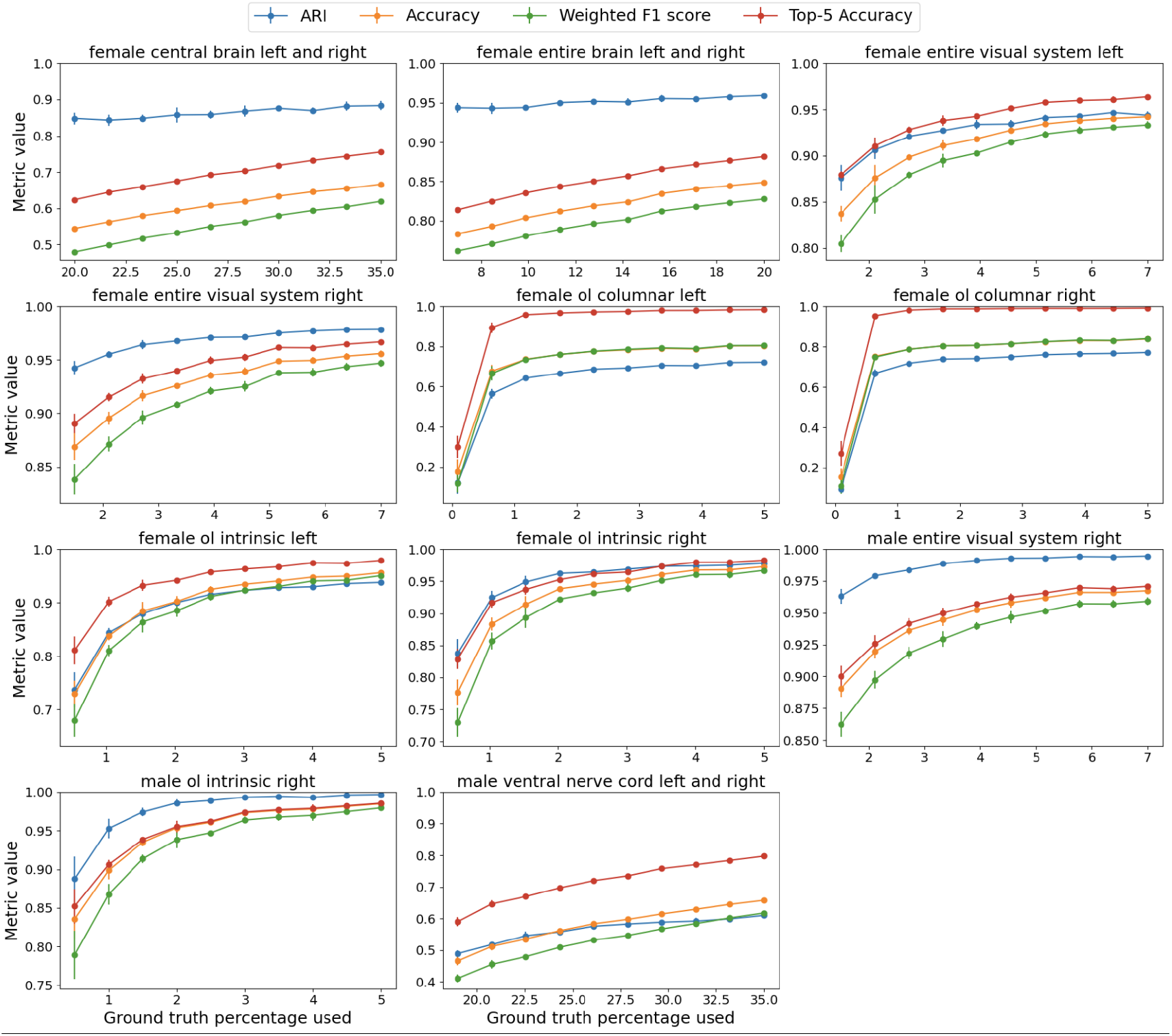
Detailed plots per dataset. We plot accuracy, top-5 accuracy, ARI and weighted F1 as a function of labeled nodes. We sample the labeled nodes uniformly at random, and do not guarantee that there is at least one labeled node per class.

Next, we list detailed per-region results. We utilize the following metrics: Accuracy, Top-5 Accuracy, Adjusted Rand Index (ARI), and weighted F1 score. Accuracy is the percentage of nodes classified correctly, top-5 accuracy is the proportion of nodes for which the correct label appears among the top 5 predicted labels, ARI measures how similar the clustering structure is to the ground truth cluster structure, adjusting for chance groupings, and weighted F1 averages the F1 scores of all classes, weighting each class’s contribution by its number of true instances. We ran our algorithm for 15 iterations and repeated every experiment 5 times with different random seeds.

While the assumption that we have a single labeled element per class is rather mild, we also provide results for simple uniform sampling. We observe that the quality in the optic lobes remains (mostly) unaffected, while it degrades in the central brain where the number of cells per class is significantly lower (see Table 1 in Supplemental Data).

### Comparison against Seeded NBLAST with k-NN

NBLAST computes a pairwise morphological similarity score between neurons. To benchmark our semi-supervised method, we first build an NBLAST similarity matrix for neurons in the right intrinsic optic lobe of the female brain (where entry (i,j) is the similarity score between neurons i and j). We then randomly sample a fraction of neurons, guaranteeing at least one per cell type, and use these as labeled inputs to build a k-Nearest Neighbor (k-NN) classifier (with distance defined as 1 – similarity score). As Fig. 7 shows, even with 35% of neurons labeled, the NBLAST-based k-NN remains below 50% accuracy on the visual system, whereas NTAC surpasses 90% accuracy with under 1% labeled neurons. On the other hand, it achieves results comparable to NTAC on the central brain. We would like to emphasize that computing the NBLAST scores accurately requires generating high resolution skeletons and is computationally intensive, taking hours or more on a dataset with 10,000+ cells.

**Figure 7.**
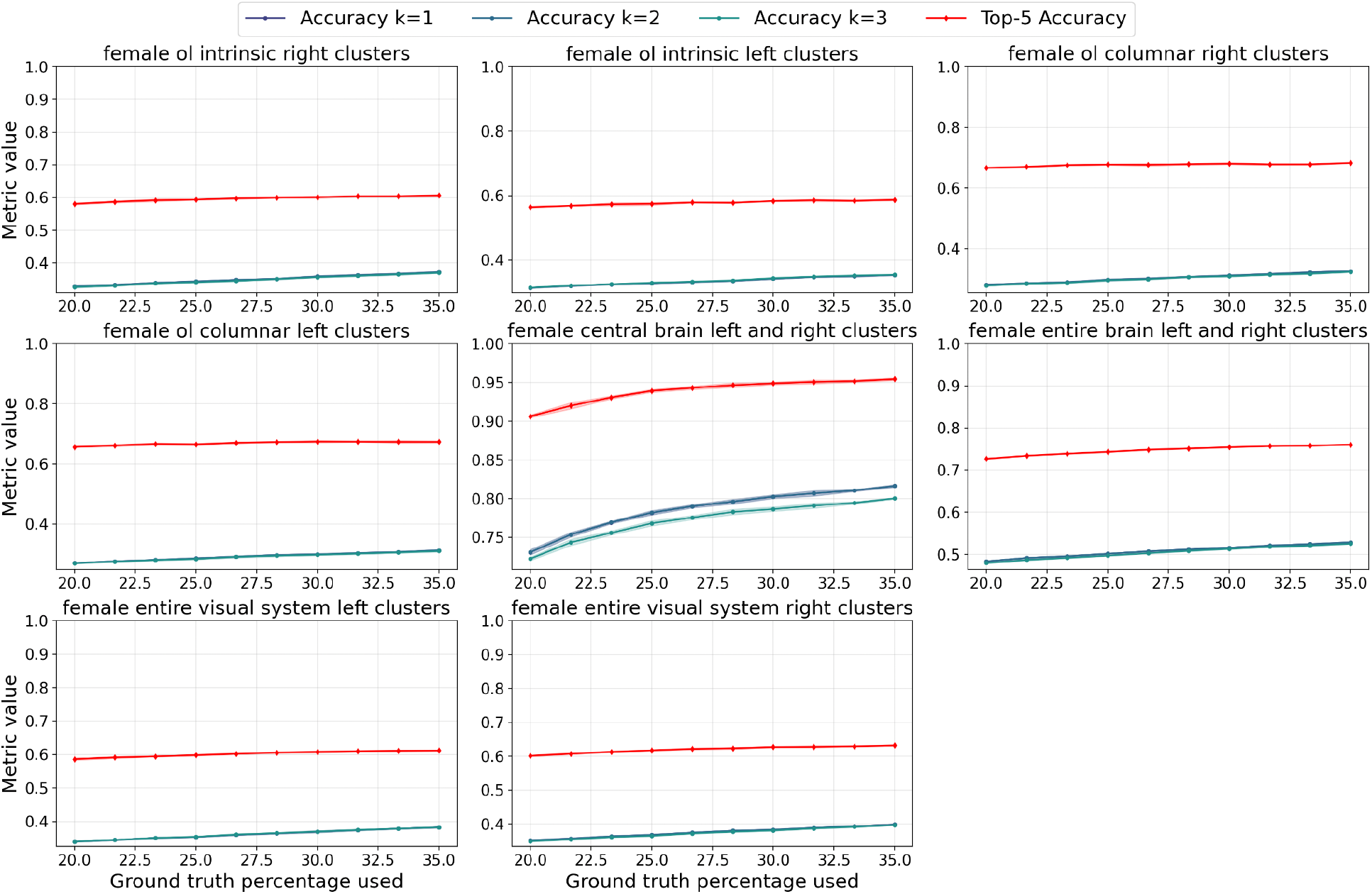
Performance of the NBLAST baseline on various regions of the female brain. This plot shows the performance of a k-Nearest Neighbor classifier using NBLAST similarity scores to infer distances as we vary the proportion of labelled neurons provided to the classifier. For k larger than 1, weighted voting is used where weights are proportional to inverse distances. For top-5 accuracy, we take the top 5 *unique* labels. Shaded regions show standard deviation over 5 repeats.

## Unseeded NTAC

### Setup

#### Parameters

The parameter *k* (maximum number of clusters) was set to the number of distinct labels (*K\**) in the ground truth. The number of iterations for the seeded subproblems was set to *R*=12. Since the computational bottleneck is selecting a new seed, we have also added an optional percentage parameter *T* which controls the maximum fraction of nodes to consider for the next seed selection, as described right after Algorithm 2.

#### Accuracy metrics

As in the previous section, we compute the following metrics: Accuracy and weighted F1 score. We also report the *Average Jaccard Cost* (our cost function *cost*(*𝒫*) divided by the number of vertices for normalization). Note that the unseeded algorithm outputs an unlabelled clustering, as labels are arbitrary. In order to compute these metrics, we need to compare the ground truth solution *G* with *K\** clusters to a candidate solution *S* with *K ≤ K\** clusters. To this end, we find a maximum matching in a bipartite graph whose nodes represent clusters in *S* and clusters in *G*, and the weight between cluster *A* from *S* and cluster *B* from *G* is given by the number of common nodes |A ∩ B|. The matching yields distinct labels for each cluster in *S*, which we then use to compute Accuracy and weighted F1 scores.

## Results

### Running time

The full running time is approximately 3.5 hours for each of the three visual systems when all vertices are considered for candidate seed selection (*T* = 100%). If instead we set *T* = 10%, each of the three visual systems ran in roughly 2 hours while attaining the same accuracy (see Fig. 8). Interestingly, in all three of them, maximum accuracy was reached within the first 30 minutes. See Methods for compute environment details.

**Figure 8.**
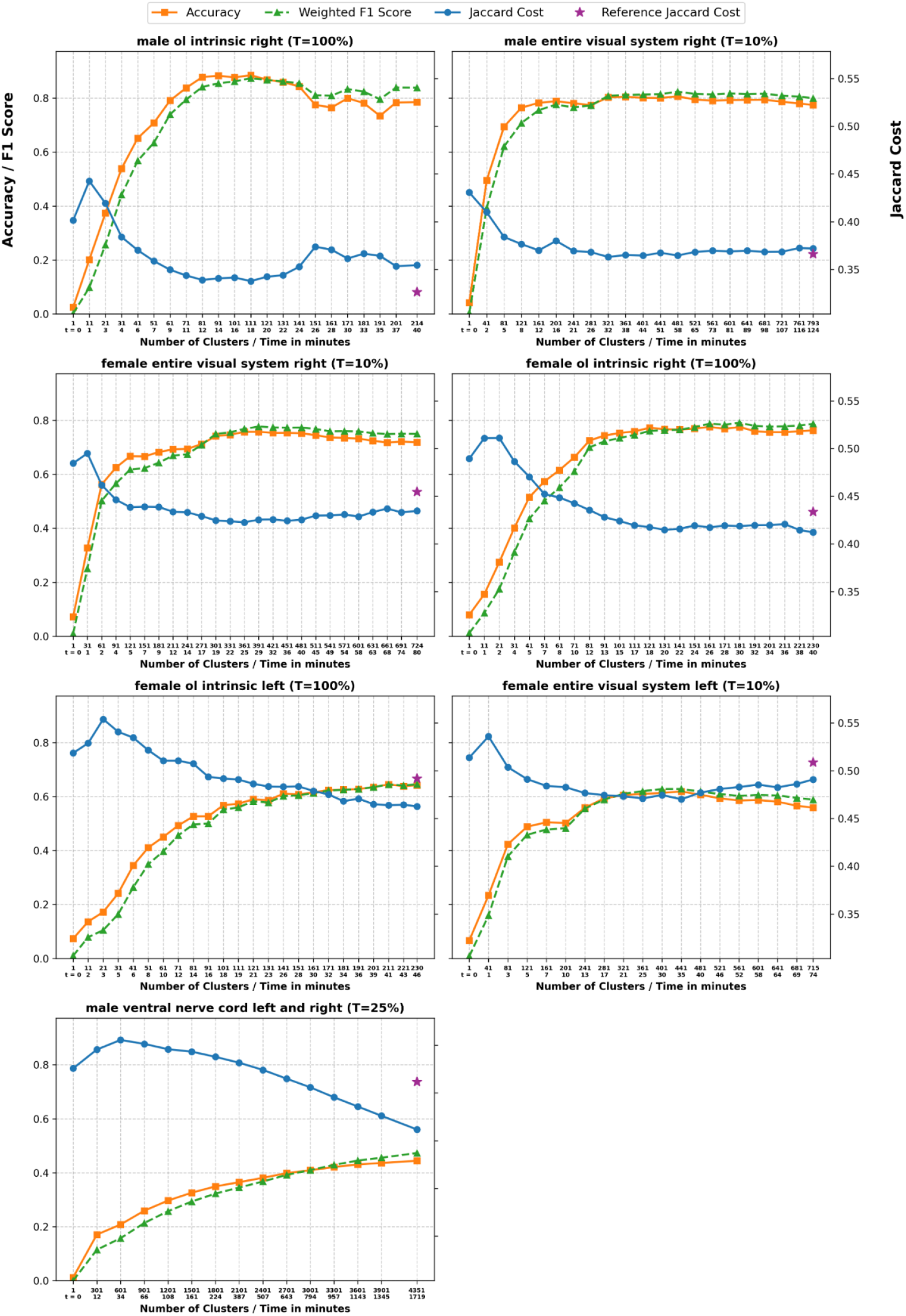
Detailed plots per dataset. We plot accuracy, weighted F1 score and Jaccard cost as a function of the number of clusters returned by the algorithm. The second line of the x-axis corresponds to the time in minutes it took the algorithm to run. T is the seed selection parameter. The star represents the Jaccard cost of the ground truth.

**Figure 9.**
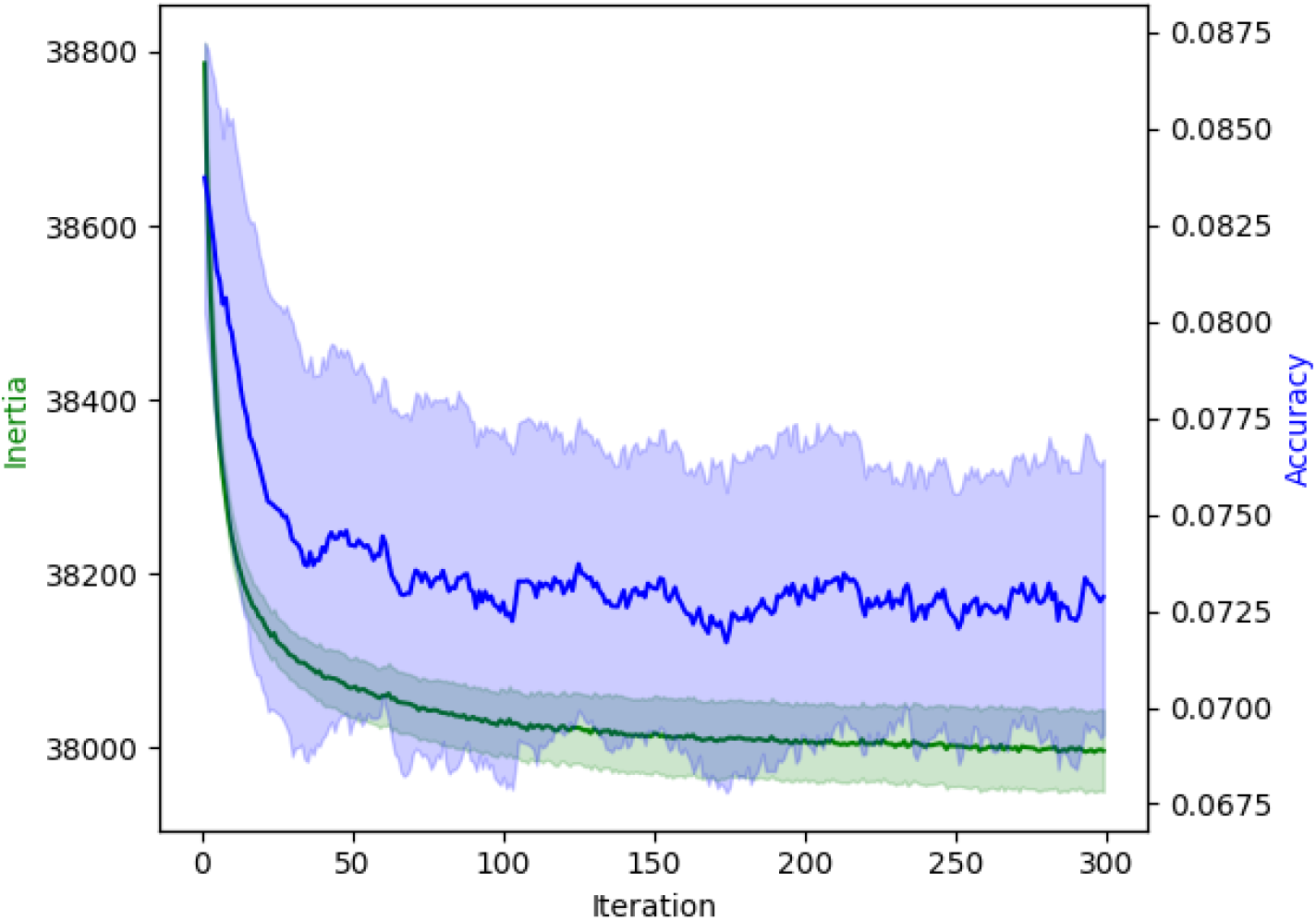
Performance of kernel k-means on the NBLAST similarity matrix of female right ol intrinsic. The x axis indicates the number of k-means iterations, the left y axis shows the objective value of the kernel k-means optimization problem (“Inertia”), and the right y axis shows the accuracy of the clustering with respect to the ground truth labels. Shaded regions show standard deviation over 5 repeats.

### Accuracy metrics

Fig. 8 shows accuracy as a function of the number of clusters/running time. As expected, we observe a negative correlation between accuracy and the Jaccard cost we are attempting to minimize. We notice that increasing the number of clusters does not always result in a better solution, which is particularly noticeable in the left female visual system plot. Furthermore, among all the partial solutions with fewer than the maximum number of clusters, the most accurate solution tracks very closely the one with the smallest Jaccard cost. For this reason, our implementation of NTAC returns both the last solution found (with the user-specified number of clusters k) and the lowest-cost solution with possibly fewer clusters, which can be more informative.

## Comparison against Unseeded NBLAST with k-NN

To compare our method against NBLAST in the unsupervised setting, we first compute a similarity matrix using the NBLAST scores on the right intrinsic optic lobe region of the female brain. The (i,j)th entry of the similarity matrix corresponds to the NBLAST score between the ith and jth neurons.

We then run kernel k-means (“Kernel K-Means,” n.d.) using this similarity matrix. As shown in Fig. 9, this method performs substantially worse than ours, failing to reach even 10% accuracy at convergence—whereas our unsupervised approach achieves 75% on the same dataset (Fig. 8). Notably, the kernel k-means objective (inertia) appears misaligned with clustering accuracy: although the algorithm consistently lowers its objective over time, this corresponds to a *decrease* in accuracy. Combined with the fact that kernel k-means is prohibitively slow on larger datasets, we conclude that it is not a viable unsupervised strategy for this problem.

## Experimental Setup

The unseeded code was executed on a system with an Intel Core i7-1355U processor with 10 physical cores and 12 logical processors, with a maximum clock speed of 5.0 GHz and 16GB RAM. For GPU acceleration, the system uses an NVIDIA GeForce RTX 2070 with 8GB VRAM running CUDA 12.4. The seeded code was executed on a MacBook Pro (Apple M2 Max, 96 GB RAM).

## Code Availability

NTAC code (Python) is open and available at github.com/BenJourdan/ntac with instructions for installation, usage, and example datasets for testing. CUDA/GPU acceleration option is available for large datasets. Contributions and feedback are welcome.

## Data Availability

The datasets used in this study have been fully proofread and annotated, and are publicly available: FlyWire full-brain connectome (https://flywire.ai), Janelia Research Campus’s *Drosophila* optic lobe connectome (https://www.janelia.org/project-team/flyem/optic-lobe) and Ventral Nerve Cord (https://www.janelia.org/project-team/flyem/manc-connectome). Snapshots of these datasets are also available for web-based exploration on the Flywire Codex (Matsliah et al. 2023): https://codex.flywire.ai/?dataset=fafb, https://codex.flywire.ai/?dataset=maol and https://codex.flywire.ai/?dataset=manc.

## Discussion and future work

Our findings indicate that connectivity-based analysis presents a compelling alternative to current state of the art neuron classification methods that rely on anatomical characteristics. This approach demonstrates greater robustness and efficiency, particularly in contexts of large scale connectome reconstructions where morphological analysis is hindered by high variability in neuronal structures and the uniqueness of individual synaptic partners—as observed in the visual systems of *Drosophila* and the mammalian visual cortex.

Future work may focus on improving classification accuracy in the unseeded regime and removing the need to predefine the maximum number of clusters (*K*). In addition, when run on partial connectomes/systems, NTAC is prone to errors on boundaries where neuronal partners are missing (see Fig. SD1). On the theoretical side, intriguing open problems are to develop provable approximation guarantees for approximate equitable partitioning, and to devise more efficient algorithms. Additionally, incorporating multimodal data—such as spatial positioning and arborization patterns—holds potential to further enhance classification performance and yield a more comprehensive understanding of neuronal cell types.

## Acknowledgements

We thank Chris Salmon, Mala Murthy, Sebastian Seung and Jakob Troidl for insightful discussions and suggestions. Arie Matsliah was supported by funding provided through grants to Murthy and Seung from the NIH BRAIN Initiative (RF1 MH117815, RF1 MH129268, U24 NS126935). Gregory Schwartzman was supported by the following research grants: KAKENHI 25K00370, JST ASPIRE JPMJAP2302 and JST CRONOS JPMJCS24K2.

## Supplemental data

### Datasets Statistics

**Table SD1.**
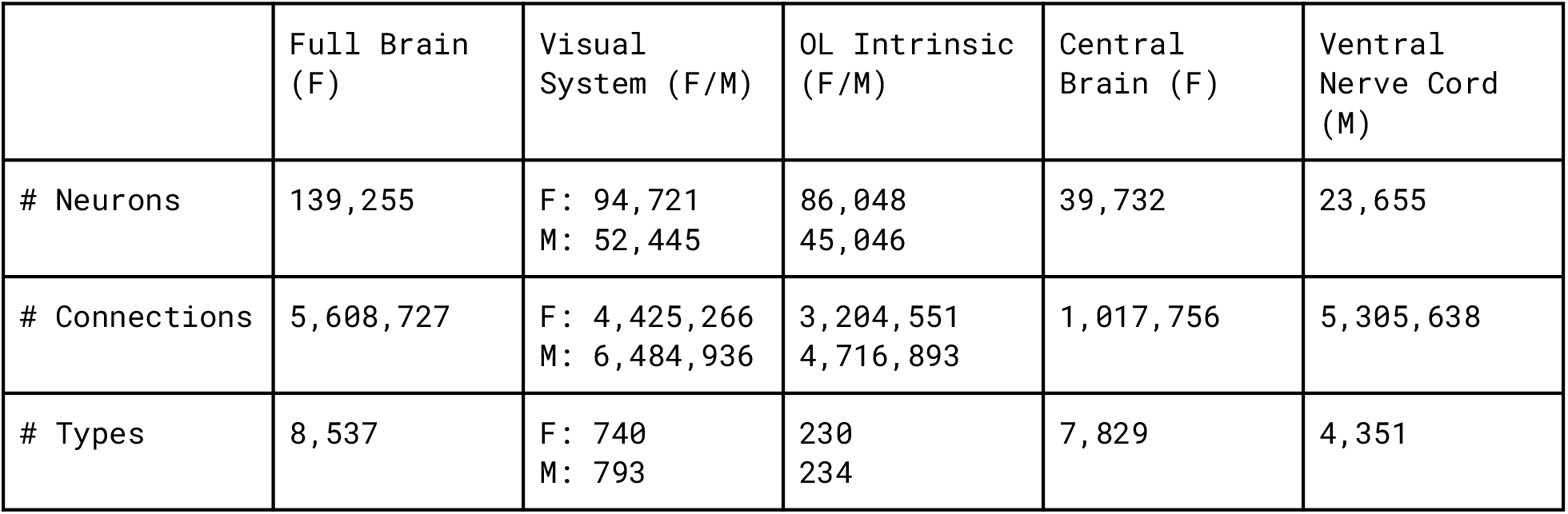
The male visual system contains only the right hand side, the female contains both. OL Intrinsic is a subset of the visual system, which itself is a subset of the full brain. Central brain is a subset of the full brain. Full brain and the ventral nerve cord together form the CNS, while neck neurons belong to both. See illustration in Fig. 1.

#### Analysis of inaccuracies

Taking a deeper look into cases where NTAC struggles on the columnar3 neurons we observe that the inaccuracies occur on the boundaries (T4*, T5* neuron types). Using 5% labeled neurons yields 80% accuracy, compared to >95% accuracy on complete visual system.

**Figure SD1.**
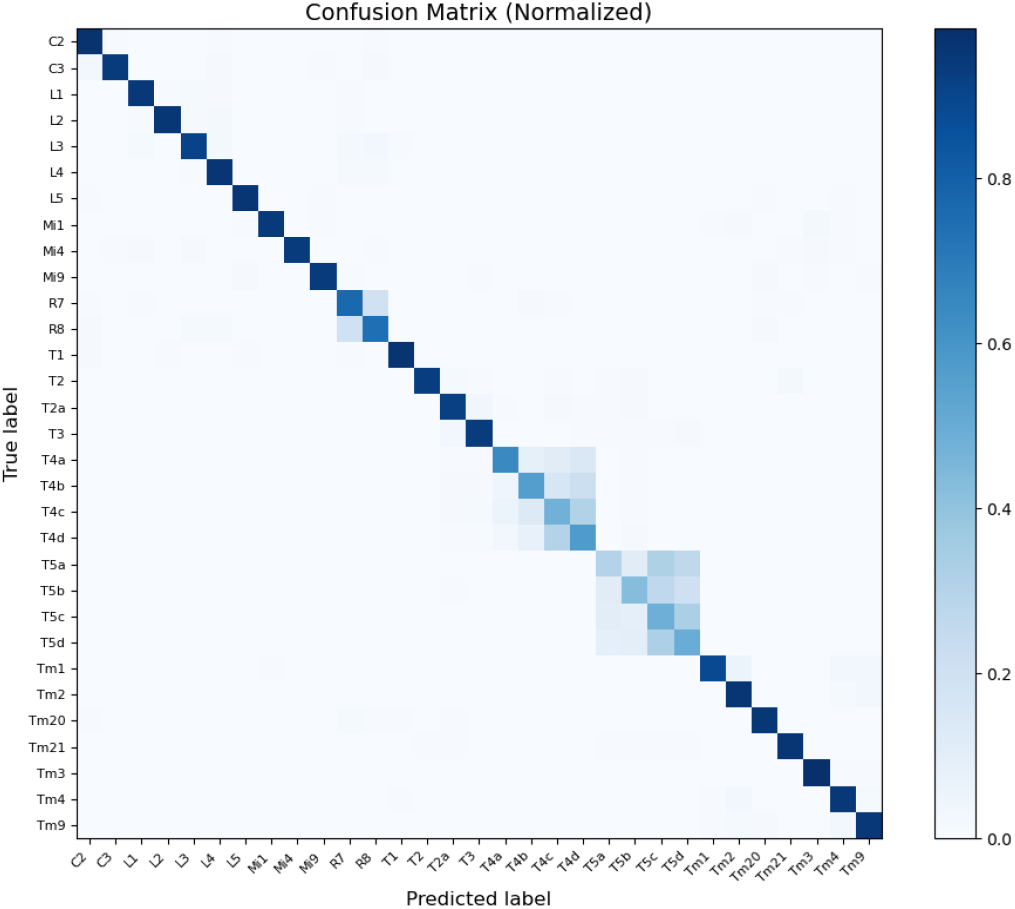
Confusion matrix for the columnar types. The most problematic classes are T4 and T5 subtypes, and they are misclassified among themselves when the dataset at hand does not include downstream projection neurons to differentiate between them.

**Figure SD2.**
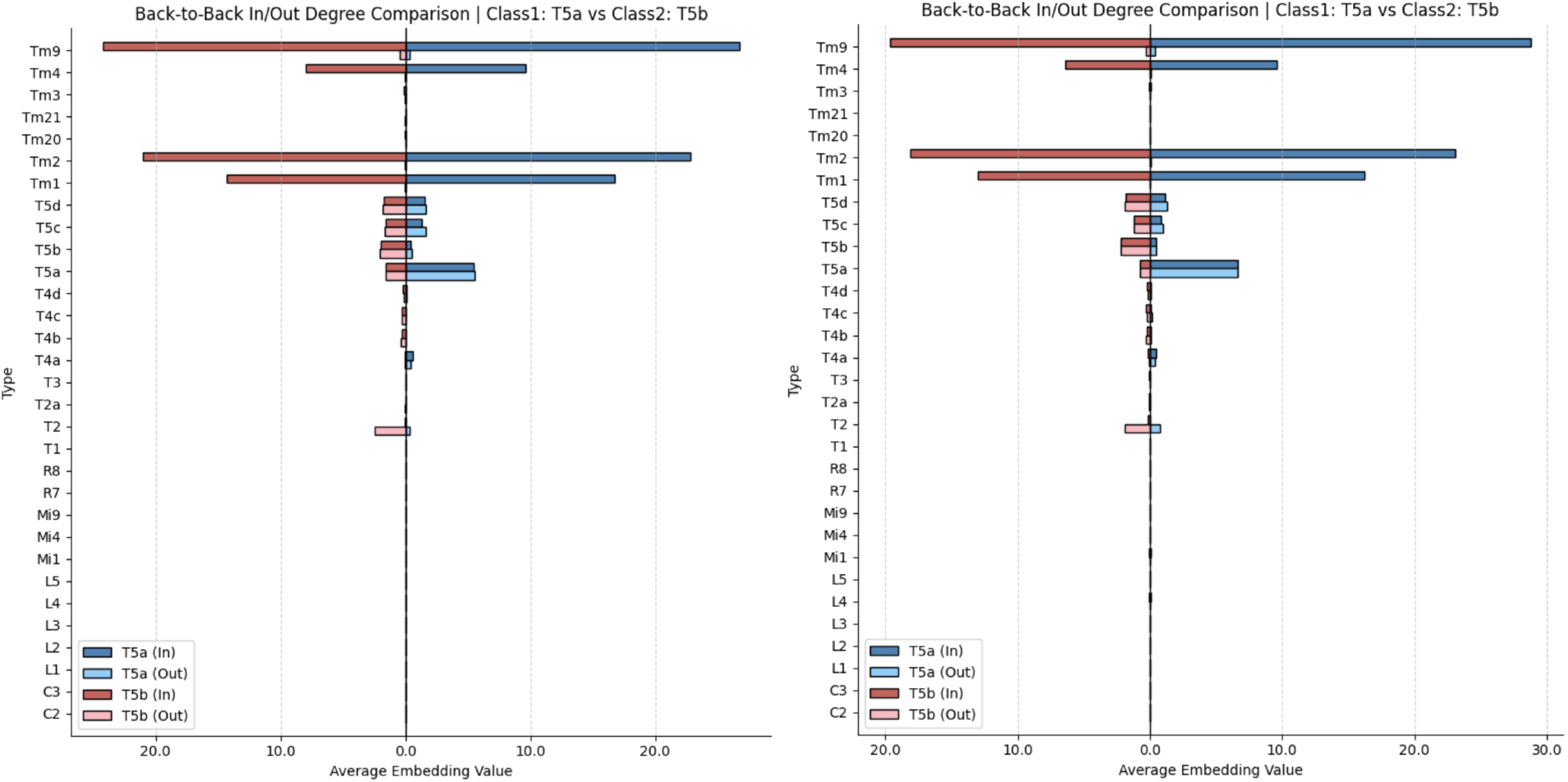
The average neighborhood histogram (the “fingerprints”) of T5a compared to T5b. On the left we use the ground truth partition and on the right we use the partition returned by NTAC. Plots show the average neighborhood value for a specific type over all neurons in the class. This implies that using only the topological features of the columnar neurons, the T5 types are nearly impossible to distinguish. However, when classifying a larger dataset (not just the columnar cells in isolation), the algorithm behaves much better because their downstream partners (visual projection neurons) carry the information to differentiate them. Indeed, running NTAC on all intrinsic neurons in the optic lobe we get near perfect accuracy (99%) on both T4 and T5 type neurons.

**Figure SD3.**
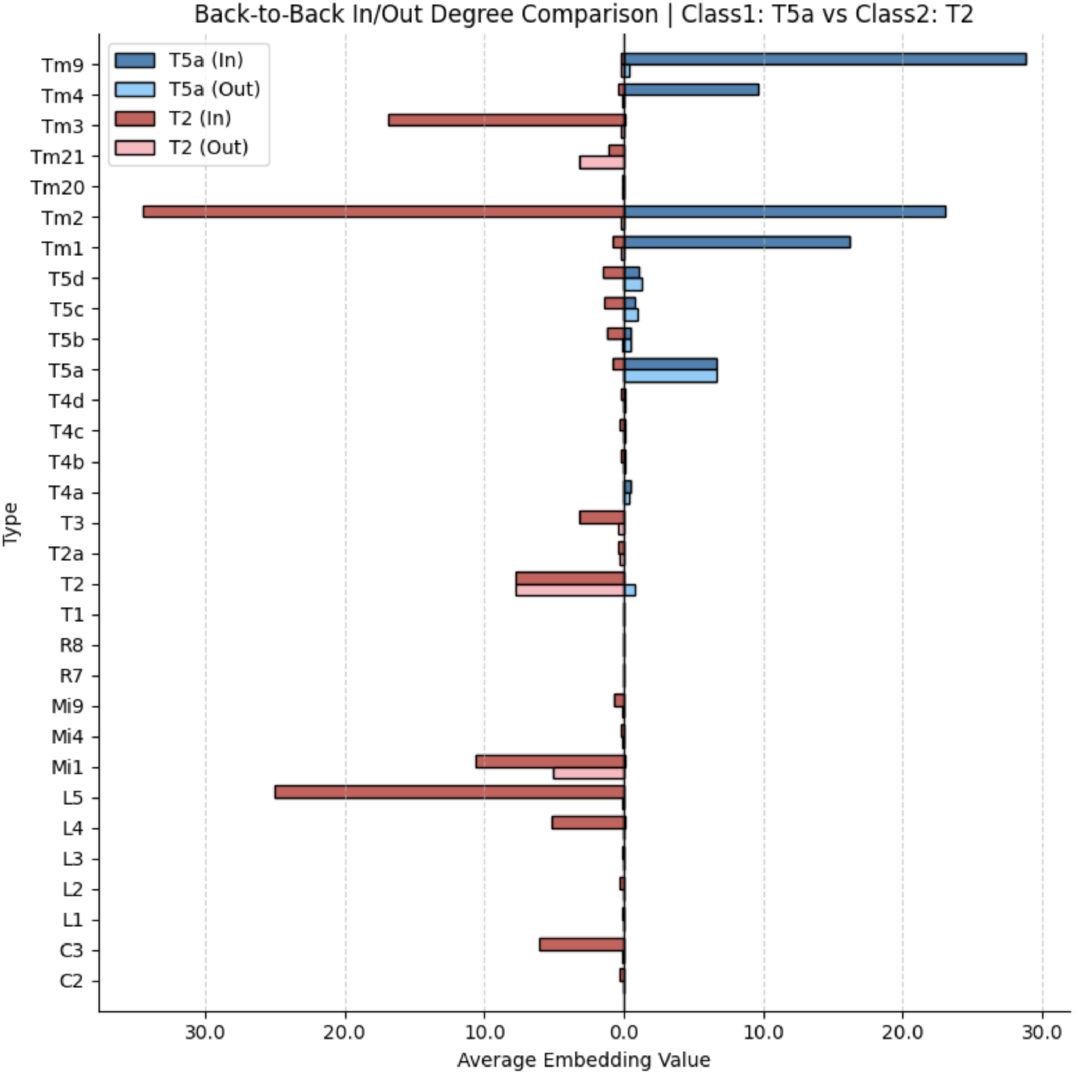
As a sanity check we compare T5a and T2: we can see that there is a significant difference.

**Figure SD4.**
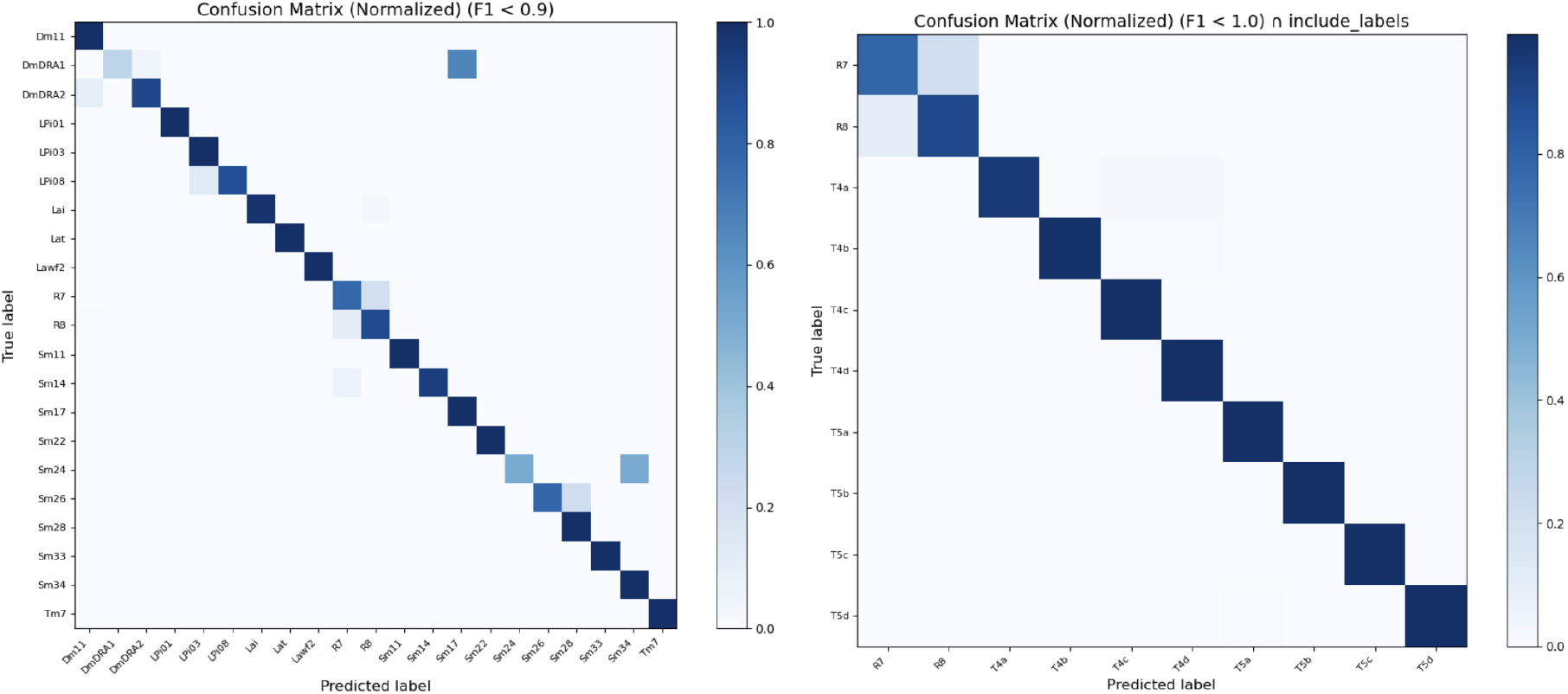
The confusion matrices for the intrinsic dataset. As the number of labels here is much larger, we plot one confusion matrix for all types which have an F-score below 0.9, and another matrix for the problematic types from the columnar dataset. We also note that the confusion between R7 and R8 persists even in the larger dataset, and this is expected since in the FlyWire connectome the reconstruction quality near the edge of the lamina was affected by a partial severing of the lamina from the medulla, which reduced the fidelity of photoreceptor reconstructions, including R1–R6 and R7/R8, and led to lower synapse attachment rates in this region (Dorkenwald et al. 2024).

**Figure SD5.**
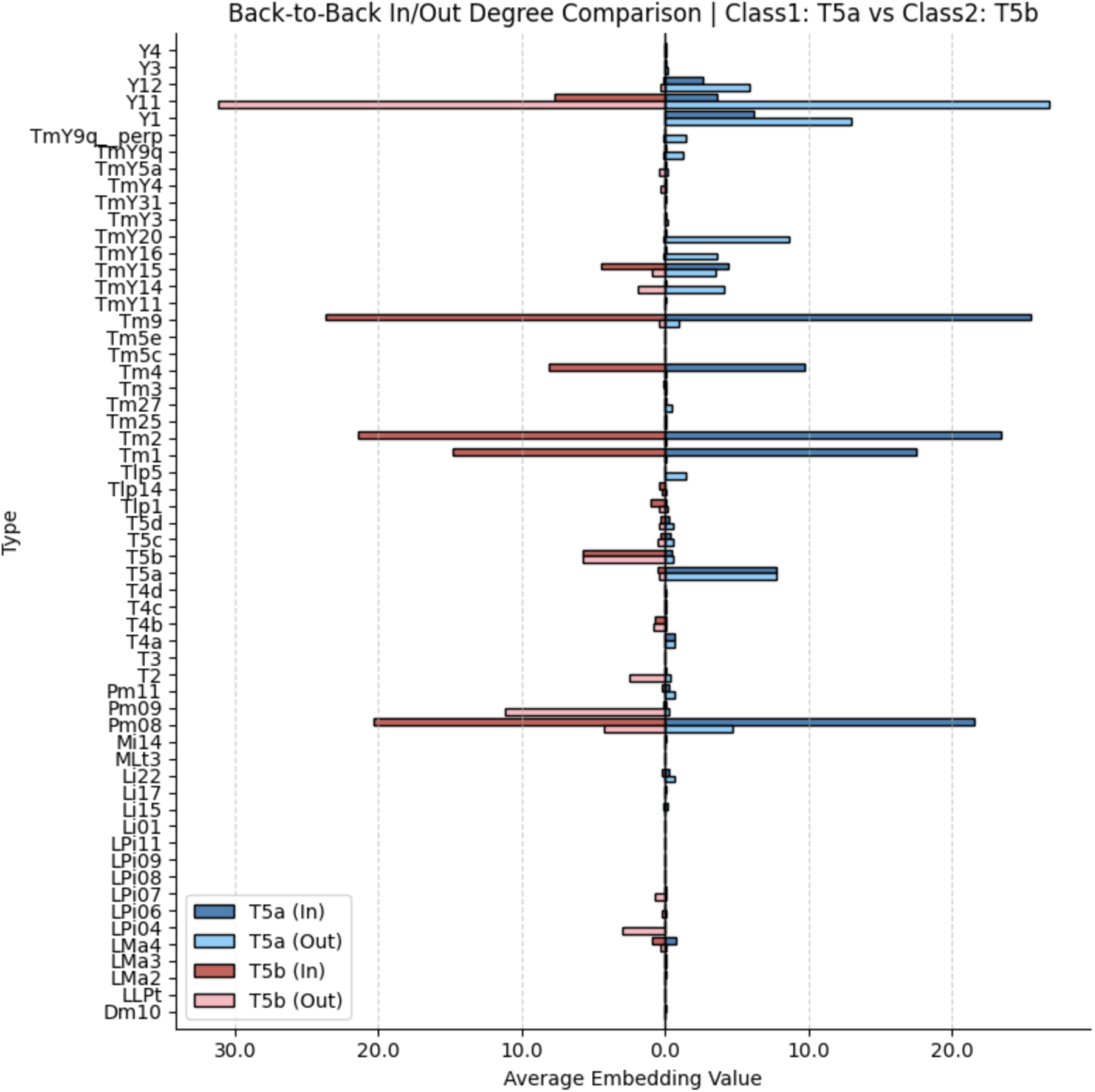
The histograms of T5a and T5b in the larger dataset. As the embeddings are very large now, let us only keep entries where both values exceed a threshold of 0.05 so that we can generate a meaningful diagram. We can now see that the two classes can be separated using topology alone.

## Footnotes

1 A neuronal cell type is a population of neurons that share similar molecular, anatomical, and physiological properties. In plain words, it is a group of brain cells that are alike in the kinds of molecules they contain, their shape/position, and their function.

2 In the *Drosophila* Central Nervous System there are roughly 160,000 neurons: 90,000 in the visual system, 40,000 in the central brain and 25,000 in the neck and nerve cord. While most neurons are in the visual system, there are only about 750 visual cell types - an order of magnitude less than in the central brain and in the nerve cord. See Table 1 in Supplemental Data.

3 Cell types in the FlyWire Optic Lobes which consist of at least 700 instances per hemisphere: C2, C3, L1, L2, L3, L4, L5, Mi1, Mi4, Mi9, T1, T2, T4a, T4b, T4c, T4d, Tm1, Tm2, Tm4, Tm9, Tm20, R7, R8, T2a, Tm3, T3, Tm21, T5c, T5b, T5a, T5d.

